# Assessment of fasting blood glucose, serum electrolyte, albumin, creatinine, urea and lipid profile among hypertensive patients and non-hypertensive participants at wolaita sodo university teaching and referral hospital, SNNPR, Ethiopia

**DOI:** 10.1101/2020.10.27.356873

**Authors:** Berhanu Haile, Mistire Wolde, Tatek Gebregziabiher

## Abstract

**Background:** Hypertension is a silent killer that requires long term management to avoid complications. It is one of major public health problem in developing counties like Ethiopia. Hypertension increases the risk of morbidity and mortality and has negative consequences on the cognitive and physical fitness of productivity in adults.

**Objective:** To assess fasting blood glucose, serum electrolyte, albumin, creatinine, urea, and lipid profile among hypertensive patients and non-hypertensive participants at wolaita sodo teaching and referral hospital.

**Methods:** A comparative cross-sectional study was conducted from December 2019 to February 2020. On the study a total of 156 study participants (78 cases and 78 controls) were involved. Each study participant, after signing informed consent, interviewed about the socio-demographic and anthropometric characteristic features. Then 5ml of the blood sample was collected from each 78 patients with hypertension and each 78 samples from apparently healthy subjects from WSUTRH during the period. Fasting blood glucose, serum electrolyte, albumin, creatinine, urea, and lipid profile level were measured in each group. The Data were analyzed by using Epi data version 3.1 and SPSS version 21.0 software (IBM Corporation, USA) and results were summarized using means and percentages and presented by using figures and tables. P-value < 0.05 was considered to be significant at 95% confidence level. Any abnormal laboratory results of study subjects dispatched and communicated with physicians for better management.

**Results:** The mean age of hypertensives and control study groups were 50 ± 10.0 and 51 ±11.3 years respectively. The body mass index of hypertensives and control study groups were 53.4% and 34.2% overweighed respectively. The mean ± SD of fasting blood glucose, total cholesterol, LDL-C, TG, RFT were significantly increases while serum sodium, calcium, albumin, and HDL-Cholesterol significantly decreased in hypertensives when compared with non-hypertensives and serum potassium was no statistical significance among case and control groups.

**Conclusion:** In present study, we observed that the hypertensive group was at risk for developing biochemical alteration in creatinine, urea, fasting blood glucose, lipid profile, electrolytes, and albumin test parameters with an increased period of time.

**Recommendation:** Regular measurements of biochemical parameters strongly needed for hypertensive patients.

## Introduction

Hypertension is higher pressure in blood vessels that harder the heart has to work in order to pump blood. If left uncontrolled, hypertension can lead to a heart attack, an enlargement of the heart and eventually heart failure. Blood vessels may develop bulges (aneurysms) and weak spots due to high pressure, making them more likely to clog and burst. The pressure in the blood vessels can also cause blood to leak out into the brain. This can cause a stroke. Hypertension can also lead to kidney failure, blindness, rupture of blood vessels and cognitive impairment (1).

Hypertension, also known as high or raised blood pressure, is a condition in which the blood vessels have persistently raised pressure. Persistent hypertension is one of the risk factors for stroke, myocardial infarction, heart failure and arterial aneurysm, and is a leading cause of chronic kidney failure. Moderate elevation of arterial blood pressure leads to shortened life expectancy. Dietary and lifestyle changes can improve blood pressure control and decrease the risk of associated health complications, although drug treatment may prove necessary in patients for whom lifestyle changes prove ineffective or insufficient. (1)

Blood pressure is created by the force of blood pushing against the walls of blood vessels (arteries) as it is pumped by the heart. Normal adult blood pressure is defined as a blood pressure of 120 mm Hg when the heart beats (systolic) and blood pressure of 80 mm Hg when the heart relaxes (diastolic). However, the cardiovascular benefits of normal blood pressure extend to lower systolic (105 mm Hg) and lower diastolic blood pressure levels (60 mm Hg). (1) Blood pressure is influenced by various genetic and lifestyle factors including nutrition. In this regard, sodium is an important mineral which, besides its functions in fluidbalance, action potential generation, digestive secretions and absorption of many nutrients, also play an important role in blood pressure regulation with a reduced sodium intake being associated with a reduction in systolic and diastolic blood pressure. Therefore, independent of body weight, sex and age, too much dietary salt (sodium chloride) is regarded as an established risk factor for hypertension. Concomitant to sodium reduction, higher potassium intake or supplementation has also been repeatedly shown to reduce the blood pressure of especially hypertensive persons. (2) High sodium and low potassium inhibit the sodium pump, increase intracellular sodium, and drive calcium into cells, which ultimately induce vascular smooth muscles contraction and increased peripheral vascular resistance. A new pathway of sodium storage in the human body has been identified. Excess sodium stored in the subcutaneous lymphatic system (on proteoglycans in interstitial space), where it becomes osmotically inactive, can act as a fluid buffering system to blunt the blood pressure increase during excessive salt intake. (3)

Hypertension is defined as a systolic blood pressure equal to or above 140 mm Hg and/or diastolic blood pressure equal to or above 90 mm Hg (1). High blood pressure has been associated with elevated atherogenic blood lipid fractions, but epidemiological surveys often give inconsistent results across population subgroups. A better understanding of the relation between blood pressure and blood lipids may provide insight into the mechanism(s) whereby hypertension is associated with increased risk of coronary heart disease. (4)

The occurrence and development of hypertension is a continuous and long-term process. Blood pressure is a sensitive index for diagnosing hypertension, can reflect the progression of hypertension to some extent. (5)

Hypertension (High blood pressure) is above 140/90 mm/Hg and it’s classified as either primary (essential) hypertension or secondary hypertension; about 90–95% of cases are categorized as “primary hypertension,” which means high blood pressure with no obvious medical cause. The remaining 5–10% of cases (Secondary hypertension) is caused by other conditions that affect the kidneys, arteries, heart or endocrine system. (6)

A review of current trends shows that the number of adults with hypertension increased from 594 million in 1975 to 1.13 billion in 2015 (7). The main modifiable causes of high blood pressure are diet, especially salt intake, obesity, excessive alcohol intake and smoking status. Globally, hypertension is responsible for 62% of cerebrovascular disease and 49% of ischaemic heart disease. High blood pressure is estimated to cause 7.1 million deaths, about 13% of the total worldwide. (8)

The aim of this study was to assess fasting blood glucose, serum electrolyte, albumin, creatinine, urea and lipid profile among hypertensive patients and non-hypertensive participants in Wolaita sodo university teaching and referral hospital.

## Methods

### Study area

The study was conducted at wolaita sodo university teaching and referral hospital which was situated at southern Ethiopia, wolaita zone, sodo town, 329 kilometers from Addis Ababa, the capital city of Ethiopia. A comparative cross-sectional study was conducted to from December 2019 to February 2020 in Wolaita sodo university teaching and referral hospital, southern national nationality people region, Ethiopia

### Sampling method and study groups

Consecutive sampling method was used to include 162 of 81 Hypertensive outpatients who visiting the Hypertensive outpatients department during the study period and 81 apparently healthy participants who were visiting the outpatient department.

### Exclusion criteria

Study groups who had history of renal disease, cardiac heart failure, pregnant women and diabetes were excluded from the study.

### Data collection

The data collection was interview with administered questionnaire mainly consisted of closed and open ended questions and delivered to eligible subjects after consent taken to collect data face to face interview, and Physical measurements obtained by trained clinical nurses.

### Whole blood sample collection and laboratory analysis

About 5 ml of venous blood was collect aseptically from the median cubital vein from each study participant by trained Laboratory Technologists in the morning after nine hours of overnight fast. The blood sample was dispense into jell coated serum separator test tube or plain tube label with unique ID number. The collected blood sample was left for 30 minutes to facilitate clotting at room temperature. Then the clotted blood samples were centrifuged for 5 minutes at 3000 revolution per minutes (rpm) to separate serum from formed elements. The fasting blood glucose, serum albumin, urea, creatinine, and lipid profiles were analyzed by Siemens Dimension EXL-200 and serum electrolyte analyzed by diamond diagnose(smartlyte) fully automated clinical chemistry analyzers are up to date with the regional and national standards. However, the serum was kept in Refrigerator at −20 °C by Nung tubes still the time of analysis greater than Seven days.

### Data analysis

all questionnaires were checked daily for completeness by the investigator and pre-coded data was entered into computer using Epi data version 3.1, and then data was transferred to SPSS version 21.0 software (IBM Corporation, USA) for further data cleaning so that to allow consistence and eliminate discrepancies, categorizing of continuous variable and finally analysis. P-value < 0.05 was considered statistically significant at 95% of Cl. Any abnormal findings of study groups reported and communicated with physician for better management.

### Ethical considerations

The study was approved by Addis Ababa University College of health sciences department of medical laboratory sciences Ethical review committee. Permission letter was written from Addis Ababa University to Wolaita Sodo University teaching and referral hospital managing director. For reasons of privacy, all data was kept confidential. Anonymity of result records were maintained by using client registration number and unique code numbers used by service providers at Wolaita Sodo University teaching and referral hospital Laboratory. The abnormal laboratory findings of study subjects were dispatched and communicated with physicians for better management.

## Results

Among the total expected162 study participants, 156 (78 hypertensive patients and 78 non-hypertensive) were included in the study with a response rate of 95.9%. Among Hypertensive patients, 40(51.3%) were females. The age ranges of Cases were from 30 −78 years old with mean age 50 ± 10.0 years. The majority of study participants 43(55.1%) from urban areas, 51(65.4%) married, 26(33.3%) no formal education, 35(44.9%) had public/private employed and 50(64.1%) had monthly income below1000 Ethiopian Birr (**Table-1**).

**Table-1:** Socio-demographic Characteristics of hypertensive patients and controls at WSUTRH, SNNPR, Ethiopia, from December 2019 to February 2020.

| Variables |  | Study Participants (n=156) |  |
| --- | --- | --- | --- |
|  |  | Hypertensive patients | Controls |
|  |  | n=78 (50%) | n= 78 (50%) |
| Age in years | <50 | 28(35.9%) | 30(38.4%) |
|  | ≥50 | 50(64.1%) | 48(61.6%) |
| Gender | Female | 40(51.3%) | 40(51.3%) |
|  | Male | 38(48.7%) | 38(48.7%) |
| Residence | Urban | 43(55.1%) | 44(56.4%) |
|  | Rural | 35(44.9%) | 34(43.6%) |
| Marital Status | Married | 51(65.4%) | 42(53.8%) |
|  | Single | 14(17.9%) | 11(14.1%) |
|  | Separated | 6(7.7%) | 10(12.8%) |
|  | Widowed/widower | 7(9.0%) | 15(19.3%) |
| Educational level | No formal education | 26(33.3%) | 22(28.2%) |
|  | Primary | 13(16.6%) | 17(21.8%) |
|  | Secondary | 16(20.5%) | 11(14.1%) |
|  | College or University | 23(29.6%) | 28(35.9%) |
| Occupation | Public /Private servant | 35(44.9%) | 34(43.6%) |
|  | Merchant | 11(14.1%) | 14(17.9%) |
|  | Farmer | 20(25.6%) | 19(24.4%) |
|  | Unemployed | 12(15.4%) | 11(14.1%) |
| Monthly income level(ETB) | ≤ 1000 | 50(64.1%) | 43(55.1%) |
|  | > 1000 | 28(35.9%) | 35(44.9%) |

Regarding lifestyle factors, anthropometric and salt dietary habits of Hypertensive patients; the majority of 70(89.9%) non-smoked, 31(39.7%) consumed alcohol, 39(53.4%) overweighed and 3(4.1%) obese (Table-2). Similarly, non-hypertensive participants; the majority of 74(94.9%) non-smoked, 17(21.8%) consumed alcohol, 25(34.2%) overweighed and 47(64.4%) normal weight (**Table-2**).

**Table-2:** Lifestyle factors, anthropometric, and salt dietary habits of study participants at WSUTRH, SNNPR, Ethiopia, from December 2019 to February 2020.

| Variables | Study Participants (n= 156) |  | P-value |
| --- | --- | --- | --- |
|  | Hypertensive group(n=78) | Non-hypertensive group(n=78) |  |
| **Smoking status** |  |  |  |
| Yes | 8(10.2%) | 4(5.1%) | 0.232 |
| No | 70(89.7%) | 74(94.9%) |  |
| **Smoking frequency** |  |  |  |
| Daily | 4(50%) | 2(50%) | 0.224 |
| Frequently | 3(37.5) | 1(25%) |  |
| Rarely | 1(12.5) | 1(25%) |  |
| **Alcohol consumption** |  |  |  |
| Yes | 31(39.7%) | 27(21.8%) | 0.511 |
| No | 47(60.3%) | 51(78.2%) |  |
| **Alcohol drinking frequency** |  |  |  |
| Daily | 9(29.0%) | 4(14.8%) | 0.587 |
| Frequently | 15(48.4%) | 6(22.2%) |  |
| Rarely | 7(22.6%) | 17(63.0%) |  |
| **Dietary salt consumption** |  |  |  |
| Yes | 27(34.6%) | 62(79.5%) | < 0.001 |
| No | 51(65.4%) | 16(20.5%) |  |
| **Physical exercises activity** |  |  |  |
| Low | 36(52.6%) | 11(14.1%) | < 0.001 |
| Moderate | 32(34.6%) | 52(66.7%) |  |
| High | 10(12.8%) | 15(19.2%) |  |
| **BMI status** |  |  |  |
| Normal weight | 31(39.8%) | 50(64.1%) | 0.015 |
| Over weight | 42(53.8%) | 27(34.6%) |  |
| Obese | 5(6.4%) | 1(1.3%) |  |

### Comparison of laboratory test parameters among study groups

Hypertensive patients were significantly increased mean ± SD of FBG, RFT, TC, LDL-C, TG (p < 0.001) respectively; while significantly lower mean ± SD serum albumin, sodium, calcium and HDL-C (p < 0.001) respectively. When compared with non-hypertensive participants and serum potassium and calcium were no statistical significance among cases and controls (**Table-3**).

**Table-3:** Comparison of laboratory test parameters among study groups at WSUTRH, SNNPR, Ethiopia, from December 201 9 to February 2020.

| Test Parameters | Study Participants (n= 156) |  | t-test for Equality of Means |  |  |
| --- | --- | --- | --- | --- | --- |
| | Hypertensives<br>(n = 78)<br>Mean ± SD | Non-hypertensives<br>(n = 78)<br>Mean ± SD | p-value | 95% CI. | |
|  |  |  |  | Lower | Upper |
| FBG (mg/dL) | 122.93 ± 9.10 | 109.68 ± 10.31 | < 0.001 | 8.336 | 14.869 |
| Albumin (g/dL) | 4.27 ± 0.59 | 5.09 ± 0.96 | < 0.001 | -1.205 | -0.661 |
| Sodium (mmol/L) | 135.82 ± 4.05 | 138.90 ± 3.23 | < 0.001 | -4.172 | -1.854 |
| Potassium(mmol/L) | 4.05 ± 0.46 | 4.12 ± 0.64 | 0.403 | -0.2494 | 0.10076 |
| Calcium(mmol/L) | 2.12 ± 0.37 | 2.24 ± 0.23 | 0.299 | -8.1561 | 2.52074 |
| Creatinine(mg/dL) | 1.14 ± 0.46 | 0.85 ± 0.15 | < 0.001 | 0.16565 | 0.40050 |
| Urea (mg/dL) | 44.65 ± 18.56 | 33.21 ± 11.43 | < 0.001 | 5.8032 | 15.3506 |
| Total Cholesterol<br>(mg/dL) | 172.10 ± 33.33 | 154.53 ± 26.24 | < 0.001 | 22.877 | 40.379 |
| HDL-C (mg/dL) | 43.36 ± 7.20 | 48.92 ± 6.54 | < 0.001 | -7.670 | -3.407 |
| LDL-C (mg/dL) | 105.55 ± 29.35 | 84.19 ± 23.29 | < 0.001 | 16.025 | 33.000 |
| Triglyceride<br>(mg/dL) | 123.19 ± 27.77 | 107.14 ± 21.48 | < 0.001 | 20.559 | 37.493 |

SD- standard deviation and student t-test was used to compare means.

### Duration of hypertension with laboratory parameters among hypertensive patients

The FBG, RFT, TC, LDL-C and TG of hypertensive patients above five years showed significantly higher when compared with hypertensive patients below five years while serum albumin, sodium, calcium, and HDL-C significantly lower observed on above five years of hypertensive patients. Serum potassium was no statistical significance within two groups (Table4

**Table-4:**
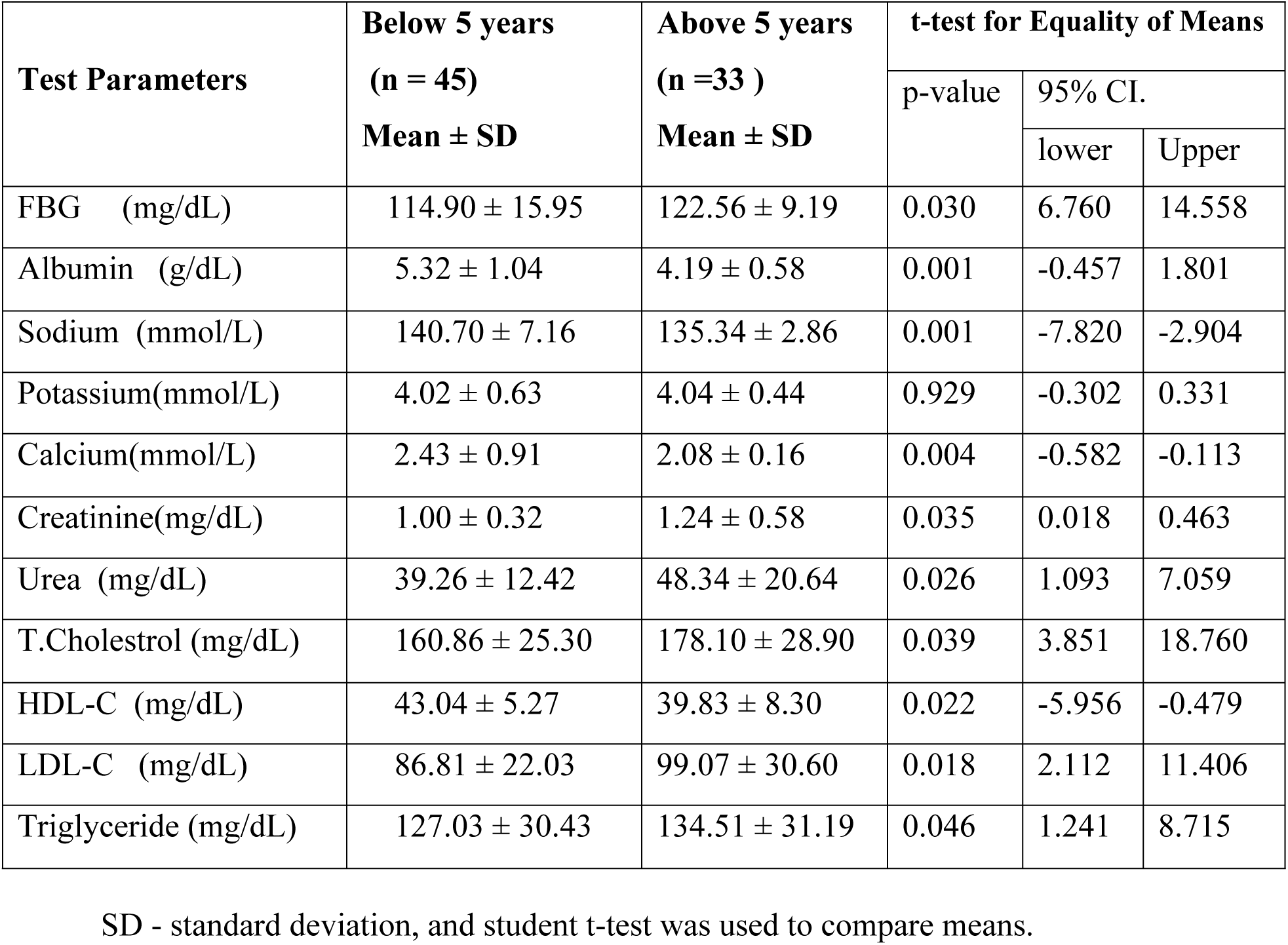
Duration of hypertension with laboratory test parameters among hypertensive patients at WSUTRH, SNNPR, Ethiopia, from December 2019 to February 2020.

| Test Parameters | Below 5 years<br>(n = 45)<br>Mean ± SD | Above 5 years<br>(n =33 )<br>Mean ± SD | t-test for Equality of Means | | |
| --- | --- | --- | --- | --- | --- |
|  |  |  | p-value | 95% CI. |  |
|  |  |  |  | lower | Upper |
| FBG (mg/dL) | 114.90 ± 15.95 | 122.56 ± 9.19 | 0.030 | 6.760 | 14.558 |
| Albumin (g/dL) | 5.32 ± 1.04 | 4.19 ± 0.58 | 0.001 | -0.457 | 1.801 |
| Sodium (mmol/L) | 140.70 ± 7.16 | 135.34 ± 2.86 | 0.001 | -7.820 | -2.904 |
| Potassium(mmol/L) | 4.02 ± 0.63 | 4.04 ± 0.44 | 0.929 | -0.302 | 0.331 |
| Calcium(mmol/L) | 2.43 ± 0.91 | 2.08 ± 0.16 | 0.004 | -0.582 | -0.113 |
| Creatinine(mg/dL) | 1.00 ± 0.32 | 1.24 ± 0.58 | 0.035 | 0.018 | 0.463 |
| Urea (mg/dL) | 39.26 ± 12.42 | 48.34 ± 20.64 | 0.026 | 1.093 | 7.059 |
| T.Cholestrol (mg/dL) | 160.86 ± 25.30 | 178.10 ± 28.90 | 0.039 | 3.851 | 18.760 |
| HDL-C (mg/dL) | 43.04 ± 5.27 | 39.83 ± 8.30 | 0.022 | -5.956 | -0.479 |
| LDL-C (mg/dL) | 86.81 ± 22.03 | 99.07 ± 30.60 | 0.018 | 2.112 | 11.406 |
| Triglyceride (mg/dL) | 127.03 ± 30.43 | 134.51 ± 31.19 | 0.046 | 1.241 | 8.715 |

SD - standard deviation, and student t-test was used to compare means.

### Prevalence of abnormal laboratory test parameters among study groups

A total of 78 hypertensive patients were included in this study, where 13(16.7%) hyperglycemia, 12(15.4%) hypoalbuminemia, 14(17.9%) hyponatremia, 8(10.3%) hyperkalemia, 15(19.2%) hypocalcemia,11(14.1%) higher creatinine, 11(14.1%) high urea, 7(8.9%) hypercholesterolemia, 5(6.4%) lower HDL-C, 8(10.3%) high LDL-C, and 10(12.8%) hypertriglycemia (**Table-5**).

Similarly, total of 78 non-hypertensive participants 6(7.7%) hyperglycaemia, 2(2.6%) hypoalbuminemia, 3(3.8%) Hyponatremia, 7(8.9%) hyperkalemia, 4(5.1%) hypocalcaemia, 2(2.6%) higher creatinine, 2(2.6%) high urea, 1(1.3%) hypercholesterolemia, 1(1.3%) lower HDL-C, 2(2.6%) high LDL-C, and 3(3.8%) hypertriglycemia (**Table-5**).

**Table-5:**
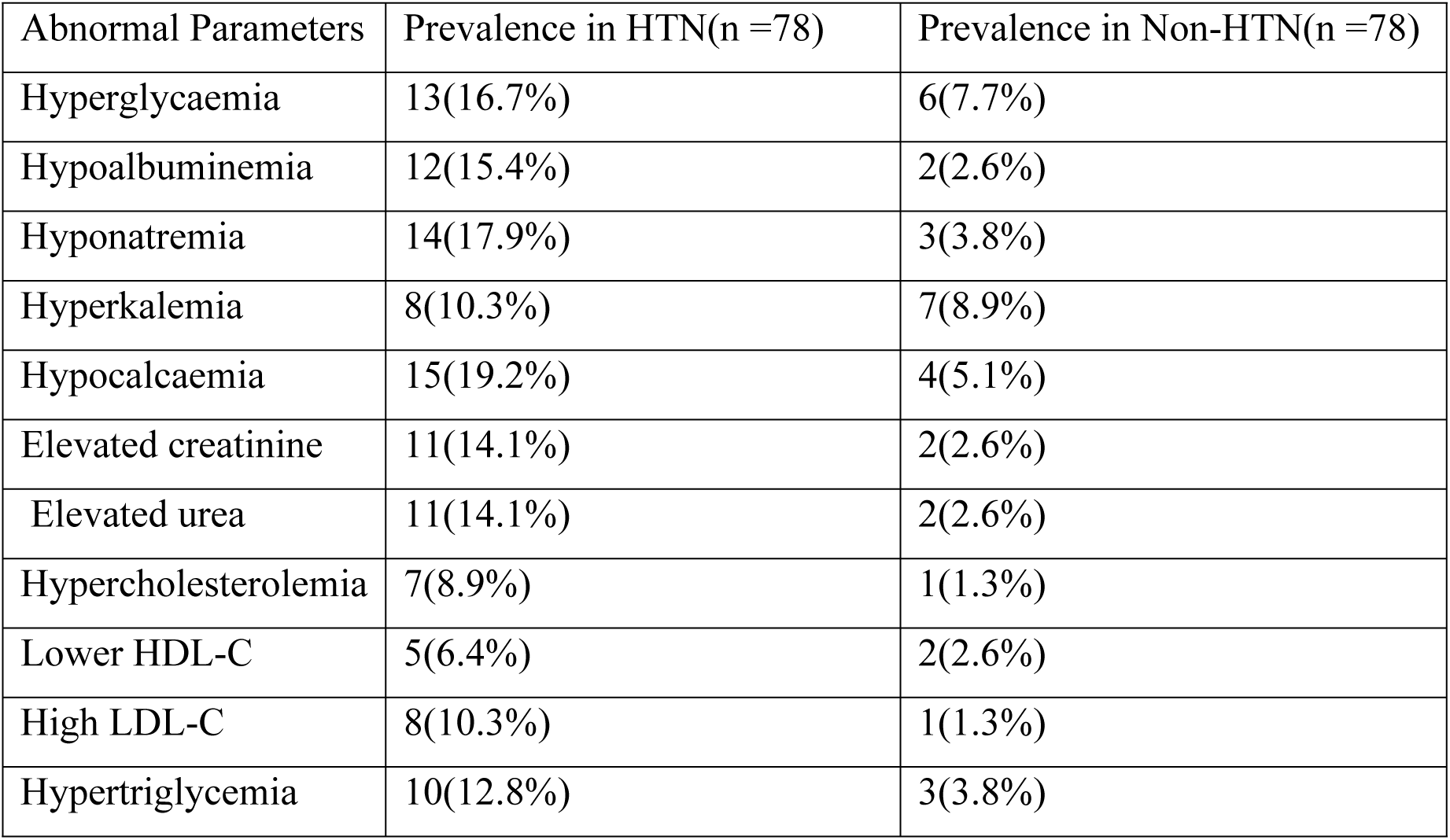
Prevalence of abnormal laboratory test parameters among study groups at WSUTRH, Ethiopia, from December 2019 to February 2020.

## Discussion

The current study assessed fasting blood glucose, serum albumin, electrolytes, creatinine, urea, and lipid profiles among hypertensive and non-hypertensive participants. The fasting blood glucose, creatinine, urea, total cholesterol, LDL-cholesterol, and triglyceride are significantly higher among hypertensive study participants when compared to the non-hypertensive study group. Sodium, calcium, albumin, and HDL-Cholesterol concentration was significantly lower among hypertensive study participants when compared to the non-hypertensive study group.

The significantly increased fasting blood glucose level of the hypertensive study participants, when compared to the non-hypertensive study group in our study, was agreed with the findings in previous studies conducted in Korean, Chinese, Cameroon, and India (9, 10, 11, 12). Hypertension was to induce microvascular dysfunction, which may contribute to the pathophysiology of diabetes development (13, 14). Endothelial dysfunction which is related to insulin resistance is also closely associated with hypertension, and biomarkers of endothelial dysfunction were found to be independent predictors of hyperglycemia. (15)

Serum creatinine and urea level were significantly higher in the hypertensive study participants when compared to the non-hypertensive study group in this study. This finding was similar to previous studies conducted in India, and Cameroon (16, 17). This may be a result of a progressing glomerular damage, endothelial dysfunction, renal microvascular disease. (20, 21) The serum albumin was significantly decreased in hypertensive study patients when compared to the non-hypertensives. This study was agreed to previous studies conducted in Japan and USA (18, 19). Hypertension is associated with endothelial dysfunction, insulin resistance, inflammation and oxidative stress, while albumin possesses both anti-inflammatory and antioxidant properties (22, 23, 24, 25). Albumin inhibits copper-stimulated peroxidation and hemolysis as well as the production of free hydroxyl radicals from systems containing copper ions and H2O2. It may also inhibit endothelial apoptosis. (26, 27)

Serum sodium and calcium were significantly decreased in hypertensive participants when compared to the non-hypertensive participants while serum potassium was no significant difference between studies groups with the findings in previous studies conducted in Nigeria (28).

The total cholesterol, LDL-cholesterol, and triglyceride were significantly elevated while HDL-Cholesterol level was lower in hypertensive study participants when compared with non-hypertensive participants with the findings in previous studies conducted in India, and Bangladesh (29, 30).

The duration of hypertension on fasting blood glucose, potassium, creatinine, urea, total cholesterol, LDL-Cholesterol, and Triglyceride concentration of hypertensive patients above five years showed significantly higher when compared with hypertensive patients for below two years. Hypertensive patients significantly lower level of serum albumin, sodium, calcium, and HDL-Cholesterol observed on the duration of hypertension occurred below two years.

Prevalence of hyperglycemia, hypoalbuminemia, hyponatremia, hypokalemia, elevated serum creatinine, elevated serum urea, hypercholesterolemia, lower HDL-C, high LDL-C, and hypertriglycemia in hypertensive patients were increased when compared with non-hypertensive subjects.

## Conclusion and Recommendation

### Conclusion

Hypertensive participants showed a significantly elevated level of fasting blood glucose, TC, TG, LDL-C, creatinine, and urea test parameters. In this study, we observed that the hypertensive group was at risk for developing biochemical abnormality in creatinine, urea, fasting blood glucose, total cholesterol, triglyceride, LDL-cholesterol, electrolytes, and albumin test parameters. The fasting blood glucose, total cholesterol, triglyceride, LDL-C, creatinine, and urea tests were significantly higher while serum albumin, sodium, calcium, HDL-C significantly decreased in the hypertensive patients with an increased period of time.

### Recommendation

- The hypertensive patients need to follow up with fasting blood glucose, serum albumin, electrolytes, lipid profile, and renal function tests for regular assessment to prevent cardiovascular disease, stroke, and kidney failure.
- Regular measurements of laboratory test parameters strongly needed for hypertensive patients.
- Further similar studies need to be conducted on hypertension that includes different study sites.

## Abbreviations and Acronyms

BMI: Body mass index
CLSI: Clinical Laboratory Standard Institute
DBP: Diastolic blood pressure
ESRD: End-stage renal disease
FBG: Fasting Blood glucose
HDL-C: High density lipoprotein cholesterol
HTN: Hypertension
IFCC: International Federation of Clinical Chemistry
ISO: International Organization for Standardization
Kg/m^2^: Kilograms per square meter
LDL-C: Low density lipoprotein cholesterol
mg/dL: mill gram per deciliter
mm Hg: millimeters of mercury
NCD: Non-communicable diseases
OPD: Outpatient department
OR: Odds ratio
RPM: Revolution per Minute
SNNPR: South Nations Nationalities and Peoples Region
SBP: Systolic blood pressure
SOP: Standard Operating Procedure
SPSS: Statistical package for social science
TC: Total cholesterol
TG: Triglyceride
WHO: World health organization
WSUTRH: Wolaita Sodo university teaching and referral hospital

## Author Contributions

Manuscript Preparation: Berhanu Haile, Mistire Wolde, Tatek G/Egzeabeher

Laboratory Analysis: Berhanu Haile

Data analysis: Berhanu Haile

Funding acquisition: Berhanu Haile

Investigation: Berhanu Haile

Methodology: Berhanu Haile, Mistire Wolde

Manuscript administration: Mistire Wolde

Writing – review & editing: Berhanu Haile, Mistire Wolde, Tatek G/Egzeabeher

## Acknowledgement

First of all thanks to God favors!!

We would thank Addis Ababa University, College of Health Sciences, and Department of Medical Laboratory Sciences for giving us this chance and cover financial fund.

Our sincerely thanks also go to the clinical Nurses who select the participants according to the selection criteria and to all participants who help us in fulfilling the questionnaire and donate biological samples.

Finally, I would like to thank for the participants to their willingness to participate and Wolaita Sodo University Teaching and Referral Hospital for all the support.

## References

1. World Health Organization. A global brief on hypertension: silent killer, global public health crisis: World Health Organization; 2013 https://apps.who.int/iris/bitstream/handle/10665/79059/WHO_DCO_WHD_2013.2_ara.pdf

2. Iqbal S, Klammer N, Ekmekcioglu C. The Effect of Electrolytes on Blood Pressure: A Brief Summary of Meta-Analyses. 2019; 11 (6):1362.

3. Frisoli TM, Schmieder RE, Grodzicki T, Messerli FH. Salt and hypertension: is salt dietary reduction worth the effort. The American journal of medicine. 2012; 125(5):433–439.

4. Bønaa KH, Thelle DS. Association between blood pressure and serum lipids in a population. The Tromsø Study. Circulation.2019; 83(4):1305–1314.

5. Lv Y, Yao Y, Ye J, Guo X, Dou J, Shen L et al. Association of Blood Pressure with Fasting Blood Glucose Levels in Northeast China: A Cross-Sectional Study. Scientific reports. 2018. https://www.nature.com/articles/s41598-018-26323-6

6. Iadecola C, Yaffe K, Biller J, Bratzke LC, Faraci FM, Gorelick PB, Gulati M, Kamel H, Knopman DS, Launer LJ, Saczynski JS. Impact of hypertension on cognitive function: a scientific statement from the American Heart Association. Hypertension. 2016; 68(6):e67–94.

7. Pereira M, Lunet N, Azevedo A, Barros H. Differences in prevalence, awareness, treatment and control of hypertension between developing and developed countries. Journal of hypertension. 2009; 27(5):963–975.

8. Weldearegawi B, Ashebir Y, Gebeye E, Gebregziabiher T, Yohannes M, Mussa S et al., Emerging chronic non-communicable diseases in rural communities of Northern Ethiopia: evidence using population-based verbal autopsy method in Kilite Awlaelo surveillance site. 2013; 28(8):891–898.

9. Scism R. Directed Evolution and Pathway Engineering for Nucleotide Analogue Biosynthesis (Doctoral dissertation, Vanderbilt University), 2010. Available at: http://etd.library.vanderbilt.edu/available/etd05112010084149/unrestricted/Scism_Dissertation.pdf

10. Kim MJ, Lim NK, Choi SJ, Park HY. Hypertension is an independent risk factor for type 2 diabetes: the Korean genome and epidemiology study. Hypertension Research. 2015; 38(11):783.

11. Cho NH, Kim KM, Choi SH, Park KS, Jang HC, Kim SS, et al. High blood pressure and its association with incident diabetes over 10 years in the Korean Genome and Epidemiology Study (KoGES). Diabetes Care. 2015; 38(7):1333–1338.

12. Yan Q, Sun D, Li X, Chen G, Zheng Q, Li L, et al. Association of blood glucose level and hypertension in Elderly Chinese Subjects: a community based study. BMC endocrine disorders. 2016; 16(1):40.

13. Feihl F, Liaudet L, Waeber B, Levy BI. Hypertension: a disease of the microcirculation? Hypertension 2006; 48: 1012–1017.

14. Nguyen TT, Wang JJ, Islam FM, Mitchell P, Tapp RJ, Zimmet PZ, et al. Retinal arteriolar narrowing predicts incidence of diabetes: the Australian Diabetes, Obesity and Lifestyle (AusDiab) Study. Diabetes 2008; 57: 536–539.

15. Meigs JB, Hu FB, Rifai N, Manson JE. Biomarkers of endothelial dysfunction and risk of type 2 diabetes mellitus. JAMA 2004; 291: 1978–198.

16. Yadav R, Bhartiya JP, Verma SK, Nandkeoliar MK. Evaluation of blood urea, creatinine and uric acid as markers of kidney functions in hypertensive patients: a prospective study. Indian Journal of Basic and Applied Medical Research. 2014; 682-689.

17. Tamanji MT, Ngwakum DA, Mbouemboue OP. Variation of Serum Uric Acid with Renal Function, Fasting Blood Glucose and Blood Pressure in Northern Cameroonians with Essential Hypertension. Annals of Medical and Health Sciences Research. 2017

18. Snipelisky D, Jentzer J, Batal O, Dardari Z, Mathier M. Serum albumin concentration as an independent prognostic indicator in patients with pulmonary arterial hypertension. Clinical cardiology. 2018; 41(6):782–787.

19. Oda E. Decreased serum albumin predicts hypertension in a Japanese health screening population. Internal Medicine. 2014; 53(7):655–660

20. Ito S, Naritomi H, Ogihara T, Shimada K, Shimamoto K, Tanaka H, et al. Impact of serum uric acid on renal function and cardiovascular events in hypertensive patients treated with losartan. Hypertension Res. 2012; 35: 867–873.

21. Ruilope LM, Salvetti A, Jamerson K, Hansson L, Warnold I, Wedel H, et al. Renal function and intensive lowering of blood pressure in hypertensive participants of the hypertension optimal treatment (HOT) Study. J Am Soc Nephrol.2001; 12: 218–225.

22. Reaven GM, Lithell H, Landsberg L. Hypertension and associated metabolic abnormalities: the role of insulin resistance and the sympathoadrenal system. N Engl J Med 1996; 334: 374–381.

23. Katagiri H, Yamada T, Oka Y. Adiposity and cardiovascular disorders: disturbance of the regulatory system consisting of humoral and neuronal signals. Circ Res 2007; 101:27–39.

24. Oda E. Metabolic syndrome: its history, mechanisms, and limitations. Acta Diabetol 2012; 49: 89–95.

25. Halliwell B. Albumin: an important extracellular antioxidant? Biochem Pharmacol 1988; 37: 569–571.

26. Zoellner H, Hofler M, Beckmann R, et al. Serum albumin is a specific inhibitor of apoptosis in human endothelial cells. J Cell Sci 1996;109: 2571–2580.

27. Yadav R, Bhartiya JP, Verma SK, Nandkeoliar MK. Evaluation of blood urea, creatinine and uric acid as markers of kidney functions in hypertensive patients: A prospective study. IJBAMR. 2014; 3: 682–689.

28. Nnadi, Henrrietta Ogadimma. Assessment of Electrolyte Levels in Hypertensive Patients in University of Port Harcourt Teaching Hospital, Port Harcourt, Rivers State, Nigeria.. American Journal of Pharm. Tech Research 2016.

29. Pyadala N, Bobbiti RR, Borugadda R, Bitinti S, Maity SN, Mallepaddi PC, et al. Assessment of lipid profile among hypertensive patients attending to a rural teaching hospital, Sangareddy. Int J Med Sci Public Health. 2016.

30. Lopes MB, Araújo LQ, Passos MT, Nishida SK, Kirsztajn GM, Cendoroglo MS, et al. Estimation of glomerular filtration rate from serum creatinine and cystatin C in octogenarians and nonagenarians. BMC nephrology. 2013; 14(1):265.

